# An snRNA-seq aging clock for the fruit fly head sheds light on sex-biased aging

**DOI:** 10.1101/2024.11.25.625273

**Authors:** Nikolai Tennant, Ananya Pavuluri, Gunjan Singh, Kaitlyn Cortez, Kate O’Connor-Giles, Erica Larschan, Ritambhara Singh

## Abstract

Although multiple high-performing epigenetic aging clocks exist, few are based directly on gene expression. Such transcriptomic aging clocks allow us to identify potential age-associated genes directly. However, most existing transcriptomic clocks model a subset of genes and are limited in their ability to predict novel biomarkers. With the growing application of single-cell sequencing, there is a need for robust single-cell transcriptomic aging clocks. Moreover, aging clocks have yet to be applied to investigate the elusive phenomenon of sex differences in aging. We introduce TimeFlies, a pan-cell-type snRNA-seq aging clock for the *Drosophila melanogaster* head. TimeFlies uses deep learning to classify the donor age of cells based on genome-wide gene expression profiles. Using explainability methods, we identified key marker genes contributing to the classification, with lncRNAs showing up as highly enriched among predicted biomarkers. lncRNA:*roX1* and lncRNA:*roX2* are top clock genes across cell types. Both are regulators of X chromosome dosage compensation, a pathway previously found to be significantly affected by aging in the mouse brain. We validated these findings experimentally in *Drosophila*, showing a decrease in survival when dosage compensation is inhibited *in vivo*. Furthermore, we trained sex-specific TimeFlies clocks and noted significant differences in model predictions and explanations between male and female clocks, suggesting that different pathways drive aging in males and females.

## Introduction

Aging is characterized by time-related dysfunction and accrued damage in an organism. Lopez-Otín et al. have suggested twelve hallmarks of aging at the molecular, cellular, and systemic levels, which underlie age-associated phenotypes [1]. A priority in the field of aging research has been the development of “aging clocks,” statistical estimators that determine the donor age of a sample based on biological measurements. These clocks allow us to discover candidate biomarkers associated with the key hallmarks of aging.

The vast majority of published aging clocks are based on DNA methylation (DNAm) data. The first aging clocks were published by Hannum et al. [2] and Horvath [3]. Hannum et al. developed an ElasticNet-based model that predicts human age from whole blood samples based on bulk DNAm levels at 71 CpG sites [2]. Horvath then developed a more robust DNAm clock, generalizable across 51 human tissue types– and even to chimpanzee tissue– utilizing 353 CpG sites [3]. A few groups have since used methylation marks to augment other clinical data points of interest in aging clock development, with the goal of understanding mortality risk and disease in the context of aging [4,5]. As many methylation marks are highly conserved, there has been an increased interest in using aging clocks to study the comparative biology of aging. Recently, the Horvath group has developed a pan-Mammalian clock that generalizes to 185 mammal species [6]. DNAm clocks exhibit high performance and have proven to be generalizable across species. The associations between DNA methylation and aging phenotypes have been widely studied for the past several decades [7,8,9,10], and the aforementioned clocks allow us to deepen our understanding.

While DNAm clocks have shown reliably high performance and have myriad contributions to various avenues of geroscience, it can be difficult to validate and apply their findings because it is often not clear which target genes are dysregulated. Epigenetic alterations, like DNA methylation, ultimately underlie changes in gene expression. DNAm aging clocks require considerable downstream analysis to determine which genes are proximal to CpG site biomarkers. Furthermore, many CpG sites identified by aging clocks are not explicitly associated with specific genes, making their significance to gene regulation events difficult to determine. Thus, transcriptomic aging clocks have the potential to reveal more direct associations between genes of interest and aging phenotypes. Identifying such genes as biomarkers of aging will provide researchers with promising targets for experimental investigation and potential therapeutic intervention, as modification of gene expression and disruption of gene products via small molecules is more feasible to implement [11].

Progress in bulk transcriptomic aging clocks has been limited due to the plethora of challenges that come with transcriptomic data. Gene fusion, alternative splicing, and post-transcriptional modifications add layers of complexity to the RNA-seq and microarray data that are difficult to disentangle. One of the first transcriptomic aging clocks fit to human peripheral blood samples obtained significant correlations between predicted and actual age, although there was a high variability across cohorts [12]. Furthermore, these clocks were trained on microarray data, a technique that has become outdated since the advent of RNA-seq due to limited dynamic range. Fleischer et al. developed a suite of regression models for an internally collected dataset of human dermal fibroblasts, which achieved noteworthy performance (r=0.81). However, this clock was not tested on external data [13]. Meyer and Schumacher found that simply binarizing RNA-seq data—that is, assigning expression values of either 0 or 1—significantly improved the performance of their *Caenorhabditis elegans* aging clock [14]. However, as gene expression exists on a continuum and is highly variable in nature, binarizing the data results in the loss of information—a binarized dataset does not properly reflect the nuances of gene expression dynamics with aging. Furthermore, the authors had to perform feature selection for an optimal set of clock genes rather than using all features in the dataset. A recent methylation clock paper showed that using all available CpG sites rather than a subset both improved model performance and created a more robust model [15], which may translate to similar results in transcriptomic clocks. Holzscheck et al. published a novel gene set-based, knowledge-primed transcriptomic aging clock using deep neural networks. This methodology yields successful performance and is highly interpretable at the pathway-level [16] but requires significant feature engineering. Genes with unknown functions would also be omitted, limiting the potential to discover new age-associated genes. Overall, while there has been progress in the development of high-performance transcriptomic aging clocks, we have yet to fully harness their potential for transcriptome-wide analysis and discovery of novel biomarkers.

Recently, there has been a rise in the popularity of single-cell sequencing because single-cell resolution unmasks the heterogeneity within biological signals, most of which are highly cell-type-specific. From a statistical and machine learning perspective, single-cell datasets have thousands of samples, thus eliminating the need to integrate several independent bulk RNA-seq datasets and address batch effects. Several single-cell aging atlases have been published, including the Tabula Muris Senis [17], the Cell Atlas of Worm Aging [18], and the Aging Fly Cell Atlas (AFCA) [19]. These data allow us to examine the dynamics of aging in different cell populations of interest. However, single-cell data poses a diverse array of computational challenges. Notably, single-cell RNA-seq often has very high dropout rates compared to bulk RNA-seq. This results in highly sparse data (a high percentage of zero values), in which the data only reflects a fraction of the cell’s gene expression. Despite these challenges, Yu et al. successfully created a single-nuclei transcriptomic clock pipeline for the aging female mouse hypothalamus. The most efficient and interpretable model, ElasticNet, which was the focus of the paper, reported an AUPRC of 0.967. However, the authors binarized the input data and subset the features to only highly variable genes rather than using the whole transcriptome [20]. Mao et al. also developed SCALE, a framework to assign a tissue-specific relative aging score at single-cell resolution to samples from the Tabula Muris Senis. However, this pipeline requires users to identify tissue-specific aging-related gene sets as input features, thus limiting the scope of novel biomarker discovery [21]. Therefore, we currently lack an interpretable single-cell transcriptomic aging clock that allows for a comprehensive transcriptome-wide analysis of aging signatures for biomarker discovery.

It is known that lifespan and healthspan are sexually dimorphic across diverse species in the animal kingdom. Yet, the innate biological mechanisms that underlie sex differences in aging remain poorly understood [22]. Sex differences in aging are seldom considered in aging research, with studies often treating sex as a confounding variable rather than a source of relevant biological variation. Understanding why aging affects males and females differently across species is fundamental to the comparative biology of aging and, on a translational level, to the development of better interventions for an aging population. As aging clocks are a framework to discover, develop, and validate hypotheses for aging biology, they can help provide insights into sex differences in aging. To our knowledge, aging clocks have yet to be used for a comprehensive study of potential genes and pathways that contribute to sex-biased aging phenotypes in any species. This leaves a crucial gap for us to begin to bridge with our investigation.

We present TimeFlies, a highly robust and accurate aging clock at single-cell resolution. We chose to develop our model based on data from the fruit fly *Drosophila melanogaster* because it is a well-studied model organism in genetics and genomics. It is an ideal candidate for studying aging and sex differences in aging due to its relatively short lifespan, extensively annotated reference transcriptome, and a plethora of widely available genetic manipulation techniques. Furthermore, the *Drosophila* brain is arguably the most well-understood across species because all the connections between individual neurons have been mapped, and individual neural circuits can be genetically manipulated [23,24]. Despite these advantages, there is a notable lack of aging clocks in the fruit fly. Thus, we chose to develop an aging clock for the *Drosophila* head to facilitate fundamental and translational studies of brain aging, especially in the context of sex differences.

We used data from the Aging Fly Cell Atlas (AFCA) [19] as input. The original AFCA paper includes aging clocks on the dataset; however, these clocks are trained on very specific cellular subtypes and have not been shown to generalize to the whole dataset, or specifically address sex differences. The highest performing clock for head tissue (r^2^ = 0.91) was specific to outer photoreceptor cells, which constitute only 7.32% of all cells in the dataset. Furthermore, the authors used regression models for these clocks, but the AFCA contains only four discrete time points across the lifespan, lending itself well to a four-class classification problem. While classifiers were also included, these were only binary classifiers for pairs of consecutive timepoints. Lastly, feature interpretation was performed on these clocks, but analysis of these features was limited to ribosomal protein-coding genes, and no analysis of sex differences in aging was included [19]. Therefore, a new clock that generalizes across all cell types in the AFCA and an exploration of sex-differential transcriptomic patterns in aging is lacking.

TimeFlies uses a 1D convolutional neural network to predict age, across the four AFCA [19] time points, from the single-cell gene expression profile. It learns from the genome-wide gene expression signals and does not require *any* feature engineering or noise reduction prior to model training. Our model generalizes across all cell types in the fly head. We have conducted an in-depth feature explainability analysis of TimeFlies for the discovery of potential aging marker genes. Our model identifies a strong role for X-chromosome dosage compensation in aging dynamics, which we demonstrate is a conserved feature between *Drosophila* and mice despite their evolutionary distance. Furthermore, we have performed sex-specific aging clock modeling to identify sex-differential transcriptomic aging signatures at single-cell resolution. Our analysis showed stark differences between male-specific and female-specific clocks, identifying pathways and functions that may be affected by aging in a sex-biased manner.

Overall, TimeFlies is a reliable aging clock based on explainable deep learning that yields valuable insights into transcriptomic aging marker discovery and opens many new avenues for future study related to sex-specific brain aging.

## Results

### TimeFlies is a pan-cell-type aging clock for biomarker discovery

TimeFlies generalizes across all cell types in the fly head with state-of-the-art performance (Test F1 score=0.9451, Test Accuracy=0.9462) in age classification (timepoints: Day 5, Day 30, Day 50, Day 70) despite very high variability between cell types (Fig 1A, “All”). However, as the dataset is not uniformly distributed across cell types, we also trained cell-type-specific clocks. We chose the five most populous broad cell types to analyze. TimeFlies maintained very high performance across all five of the cell types of interest, with the lowest performance on epithelial cells, although the F1 score was still above 0.93 (Fig 1a). Using broader cell-type categories ensured that even the cell-type-specific clocks were highly robust and could generalize across many specific subtypes.

**Figure 1.**
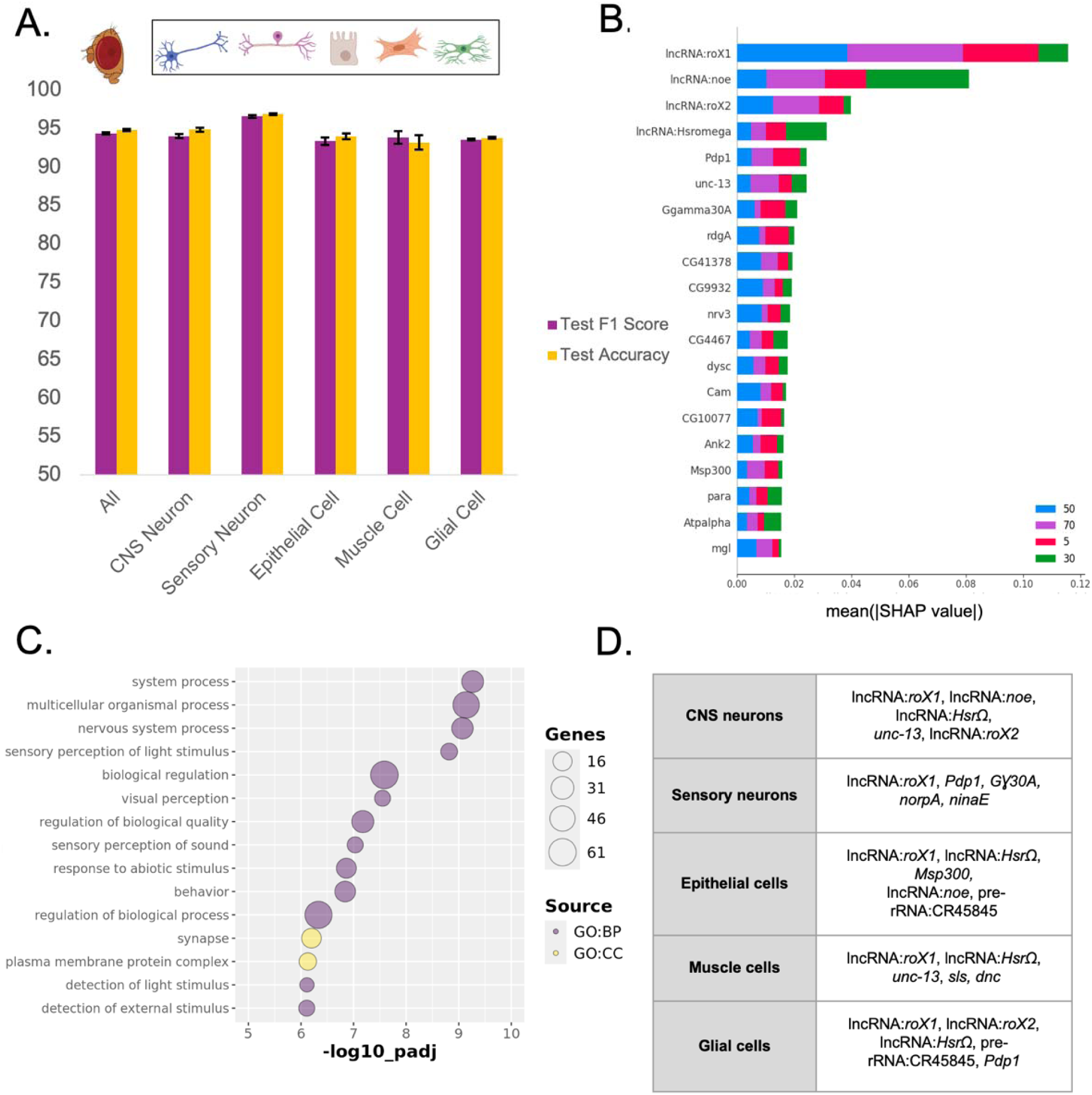
**A)** TimeFlies age classification performance on held-out test data across five cell types, across five randomly selected seeds. Error bars indicate standard deviation. **B)** SHAP summary plot showing the list of top 20 features used by TimeFlies in the classification task. Bars signify the average impact of the feature on model output magnitude. **C)** Gene set enrichment analysis performed on the top 100 genes (based on SHAP value magnitude) for the pan-cell-type clock. **D)** The top 5 genes of each cell type-specific clock, in descending order of mean SHAP value magnitude.

**Figure 2.**
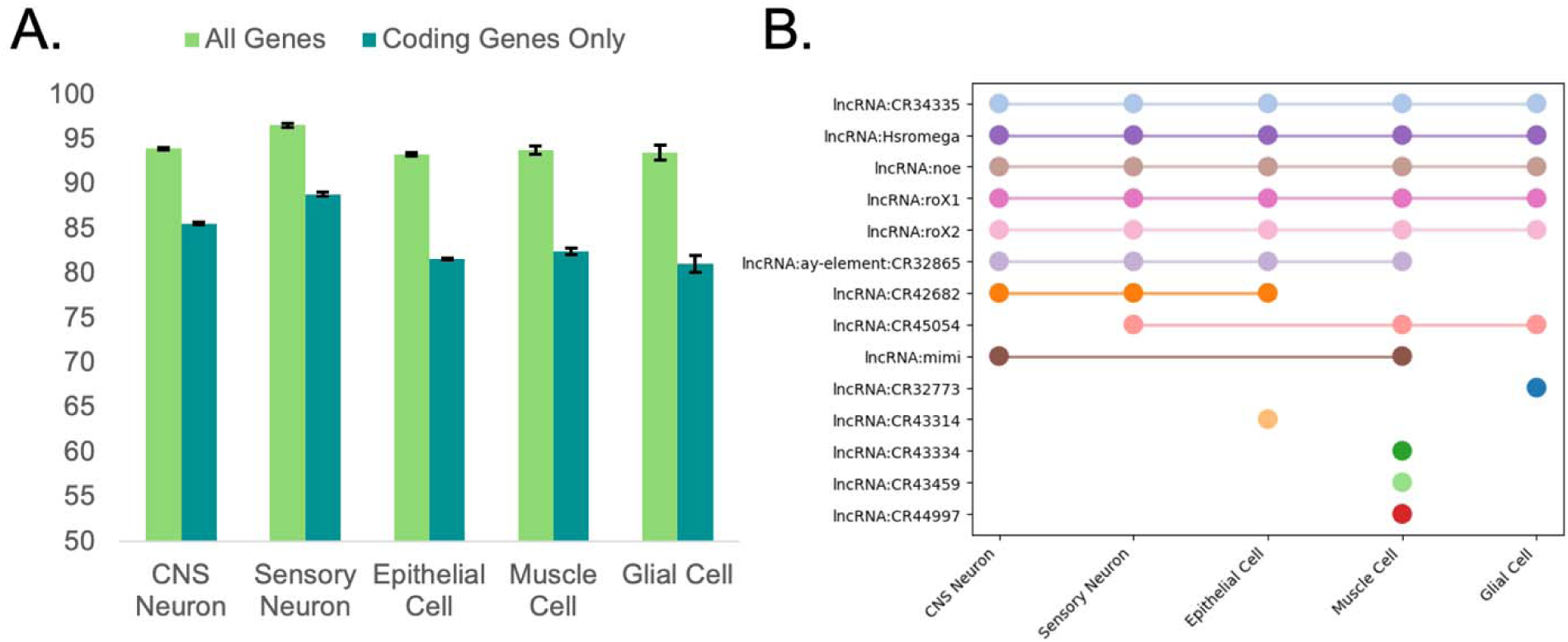
**A)** Comparison of the performance of TimeFlies (via test F1 score, y-axis) when using the whole transcriptome versus only the coding transcriptome. Average test F1 score across five random seeds is plotted with standard deviation error bars. **B)** UpSet plot of lncRNA biomarkers across cell types.

Next, we performed model explainability on the pan-cell-type clock. We determined the Shapley (SHAP) value of each gene in the dataset, which quantifies the contribution of each feature to the difference between the model’s prediction and the average base output. For a representative subset of samples from the held-out test set, SHAP values are computed for each feature for individual predictions, and averaged across all predictions to determine a global feature explanation [61]. Genes with relatively larger average SHAP value magnitudes are interpreted to be the key drivers in model predictions. We generated a SHAP summary plot (Fig 1B) of the top features from the pan-cell-type model, which shows the top 20 genes driving model predictions and a bar plot display of their average SHAP value magnitudes. We have kept the summary plot to only the top 20 genes for brevity, but have included the top 100 clock genes in Supplementary File 1. Subsequently, we ran a gene set enrichment analysis (GSEA) on the top 100 genes with the highest SHAP values (Fig 1C), which showed that many of the genes are involved in the perception of visual light (Fig. 1C). Furthermore, we determined the top features of each of the cell-type-specific clocks. TimeFlies was able to learn unique features for different cell types that were representative of each cell’s biological function (Fig. 1D, Supplementary Fig 1).

Remarkably, most features selected by TimeFlies are not among the top 5000 highly variable genes (HVGs) (Supplementary Figure 2B). However, using the whole feature set significantly outperforms using only the top 5000 HVGs in every cell type (Supplementary Figure 2A). Thus, many genes may be overlooked if only using classical differential expression analysis or performing feature engineering for simpler models. Expression patterns of *roX1*, *noe*, and *roX2* do not show significant linear associations with age due to high variability in expression levels (Supplementary Figure 2C-H). This suggests that TimeFlies detects complex age-associated patterns of expression. Thus, the explainability analysis of TimeFlies offers a more comprehensive biomarker discovery strategy than simple linear models or differential expression analyses.

The four features that were among the most significant for the pan-cell-type clock were long non-coding RNAs: lncRNA:*roX1*, lncRNA:*noe*, and lncRNA:*roX2,* and lncRNA:*Hsr*_ω_ (Fig 1C). Previous studies have identified several age-associated lncRNAs [25] and their evolutionary conservation across species [26]. It has also been shown that, in the fruit fly, lncRNAs and their known targets are differentially expressed during dietary restriction, a well-studied aging intervention [27]. Our feature explainability analysis reflects the age-associated enrichment of lncRNAs and makes the case for further evaluation of lncRNA-based gene regulation during aging.

### Noncoding RNAs are crucial to the performance of TimeFlies

After observing the prevalence of lncRNAs among the top TimeFlies genes, a phenomenon seen across cell types (Figure 1D), we sought to determine how removal of noncoding RNAs from the feature set would affect TimeFlies performance. We removed a total of 1758 noncoding genes from the dataset, with 14,234 coding genes remaining in the feature set. We re-ran cell-type-specific TimeFlies clocks. There was a drastic decrease in classification performance, assessed via test F1 score, when noncoding RNAs were omitted. This suggests that noncoding genes contribute significantly to cellular aging signatures in the fruit fly head. We then created an UpSet plot to determine which lncRNAs are common TimeFlies biomarkers across cell types. lncRNA:*roX1*, lncRNA:*roX2*, lncRNA:*noe*, lncRNA:*Hsr*_ω_, and lncRNA:*CR34335* all showed high relative SHAP value magnitudes in every cell-type-specific clock.

The lncRNA:*roX1* and lncRNA:*roX2* genes are long noncoding RNAs encoded on the X chromosome and involved in dosage compensation. This highly conserved process equalizes the levels of X-linked genes between male (XY) and female (XX) organisms. In *Drosophila melanogaster*, this process is achieved solely by upregulation of the male X chromosome, while in humans, rodents, and other mammals, one of the two female X chromosomes is silenced, which is referred to as X chromosome inactivation (XCI) prior to upregulation of the remaining X chromosome to equalize expression to autosomes [28]. Thus, X chromosome upregulation is a highly conserved process across species, including in mammals, to tune X-linked gene expression levels throughout development [29]. *roX1* and *roX2* are essential components of the male-specific lethal (MSL) complex, which facilitates hyperacetylation of H4K16 along the X chromosome targets in males, a modification associated with X chromosome upregulation. The *roX* RNAs help localize the MSL complex to the X chromosome [30, 31]. Despite the differences in structure and length of *roX1* and *roX2*, they have redundant functions due to the presence of a similar stem loop region [32]. Fascinatingly, in a single-nuclei RNA-seq study of the aging female mouse hypothalamus, *Xist*, the master regulator of X Chromosome Inactivation and the mouse analog of the *roX* genes, was the top feature in an X chromosome-based aging clock of neurons. The authors also showed that *Xist* expression is upregulated with age in some neuronal populations [33]. This suggests that, despite the evolutionary distance between mice and fruit flies, dosage compensation appears to be conserved as a significant component of the aging process.

Equally intriguing is the selection of lncRNA:*Hsr*_ω_ *(hsr*_ω_) by the model, a developmentally active gene which is inducible by heat and stress. This gene produces both nuclear and cytoplasmic transcripts [34]. The nuclear *hsr*_ω_ transcripts are essential for the organization of omega speckles, which are nuclear compartments that contain RNA binding and processing proteins. Following cellular stress such as heat shock, omega speckles rapidly disappear, with the released RNA binding proteins clustering at the *hsr*_ω_ locus, otherwise known as the *Drosophila* “93D puff.” [34, 35]. Misexpression of *hsr*_ω_ is developmentally detrimental and results in a high incidence of larval and pupal death. Downregulation of *hsr*_ω_ resulted in greater mortality in female flies than male, while overexpression of the lncRNA resulted in greater mortality of males than females [36]. Moreover, misexpression of *hsr*_ω_ is implicated in neurodegenerative disorders such as amyotrophic lateral sclerosis (ALS) and polyQ expansion disorders [37]. However, the role of *hsr*_ω_ in baseline sex-specific aging has yet to be studied, to our knowledge. Davie and colleagues published a single-cell atlas of the aging *Drosophila* brain and trained a Random Forest model to predict cellular age from gene expression values, in which *hsr*_ω_ was one of the six most influential genes [38], consistent with our analysis. Our results from TimeFlies further strengthen the importance of investigating the role of *hsr*_ω_ in aging.

The two other lncRNAs (*noe* and *CR34335*) selected by the model as biomarkers of aging across cell types both have unknown functions. lncRNA:*noe* was discovered by Kim et al. in 1998 [39]. It is abundantly expressed in the central nervous system and encodes a small peptide of 74 amino acids [39]. However, the function of the noncoding RNA and its peptide product has remained unknown since their discovery. *noe* is located within an intron of the *blot* gene [40], which encodes a sodium/chloride-dependent neurotransmitter transporter [41]. Notably, expression of *noe* is highly enriched in adult males, with moderate expression in pupae and adult females [42]. lncRNA:*CR34335* is located on the X chromosome within a long intron of the DIP-〈 gene, which encodes a neuronal cell adhesion molecule [43]. Davie et al. found that glia express high levels of lncRNA:*CR34335* while neurons express high levels of lncRNA:*noe*. Furthermore, lncRNA:*CR34335* was also among the six most influential genes in their pan-cell-type Random Forest age predictor [38], consistent with our TimeFlies results. A recently published study on wing disc regeneration found that *CR34335* localizes to the cytoplasm [44], however, the localization of *CR34335* may vary based on tissue and therefore, be different in the brain. Overall, our results suggest that both characterized and uncharacterized lncRNAs in the *Drosophila* genome may be key players in modulating age-associated pathways and thus are worth investigating.

### Sex differences in predictive aging genes

Female fruit flies, on average, have longer lifespans than male fruit flies. Thus, we developed male-specific and female-specific TimeFlies clocks to investigate sex-specific aging biomarkers. To assess the differences in what each model learned, we performed Shapley analysis and compared the model explanations of the female clock and the male clock. Remarkably, for all cell types, there were notable differences in the top clock genes for females versus males (Fig. 3A-B). LncRNAs are prevalent among the top 5 cell-type-specific clock genes for both sexes (Fig 3A-B). As expected, the *roX* RNAs are top features for all cell types in male clocks, but not the female clocks, validating that our models are able to learn sex-specific information. We then extracted the top 100 features from the female-specific and male-specific clocks for each cell type and compared the gene sets to determine the overlap of features learned by female-specific and male-specific clocks (Fig 3C). For each cell type, only 43 to 58 of the top genes from the male clock and female clock overlapped (Fig 3C). This suggests that aging, even at the cellular level, has highly sex-specific transcriptomic signatures.

**Figure 3.**
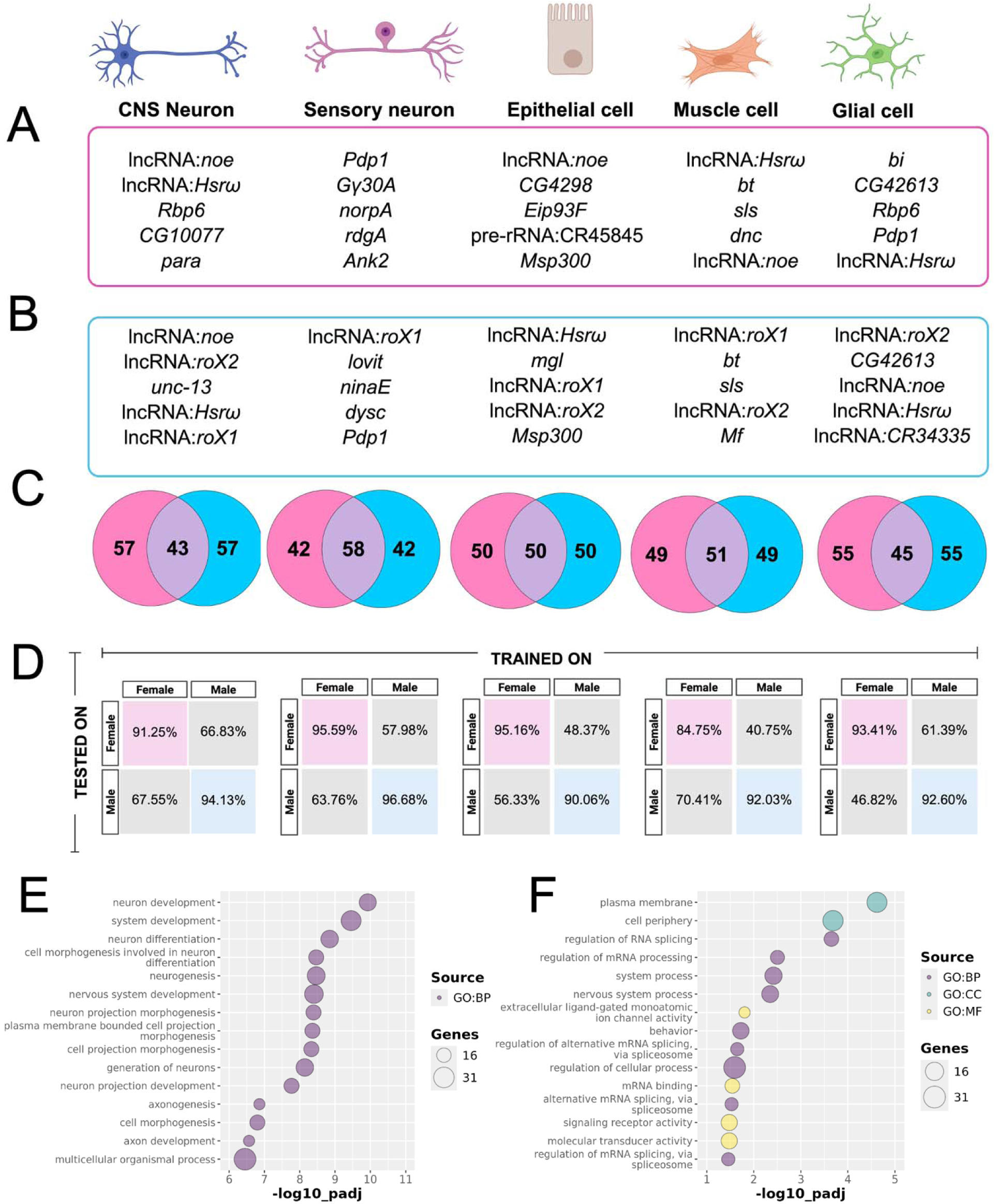
**A)** Top 5 most influential genes in age classification in TimeFlies female-specific CNS, sensory neuron, epithelial cell, muscle cell, and glial cell clocks (left to right). **B)** Top 5 most influential genes in age classification in TimeFlies male-specific CNS, sensory neuron, epithelial cell, muscle cell, and glial cell clocks (left to right). **C)** Venn diagrams showing how many genes overlapped between sex-specific clocks for each corresponding cell type; CNS, sensory neuron, epithelial cell, muscle cell, and glial cell (left to right). Pink indicates uniqueness to female and blue indicates uniqueness to male. **D)** Test F1 scores for same-sex and cross-sex training/testing in each cell type. **C)** Confusion matrix for the model trained on male samples and tested on female samples. Pink indicates trained and tested on female, blue indicates trained and tested on male, and gray indicates trained on one sex and tested on the opposite sex. **E)** GO bubble plot for genes unique to female CNS clock**. F)** GO bubble plot for genes unique to male CNS clock.

To further investigate the differences between what the male and female clocks are learning, we performed GO on the gene sets that were only found in female clocks or only found in male clocks for each cell type. For most cell types, this analysis did not elucidate any potential sex differences in aging pathways, perhaps because many of the noncoding RNAs among the sets of genes have uncharacterized function. However, the male CNS neuron clock (Fig 3F) showed enrichment of splicing-related processes, functions, and cellular components, which was not observed in the female CNS neuron clock (Fig. 3E). Regulation of alternative splicing at specific loci is known to change in baseline aging and age-associated disease in humans and mammalian model organisms [45, 46]; however, there is a lack of literature on the sex-specificity of these phenomena. In flies, sex-specific splicing has been studied throughout development, especially in the nervous system [47], but remains underexplored in aging.

We next tested whether a clock trained on only female data can generalize to male data, and vice versa (Fig 3D). Notably, clocks trained on only female samples have low performance on male samples, and clocks trained on only male samples have low performance on female samples (Fig 3D). The best results are in muscle cells and glial cells, where female-trained muscle cell clocks had an F1 score of only 40.75% when tested on male muscle cells, and male-trained glial cell clocks had an F1 score of only 46.82% when tested on female glial cells. We generated confusion matrices to determine the sources of low performance (Supplementary Fig 3). Notably, all cross-sex-tested clocks were able to distinguish 5-day-old cells with high accuracy. The male-trained clocks, apart from epithelial cell clocks, were also able to correctly classify 70-day-old female cells (Supp. Fig. 3), but most cross-sex-tested clocks struggled with the middle time points (days 30 and 50). The female-trained clocks were largely efficient in correctly classifying 50-day-old male cells, except for epithelial cells and CNS neurons (Supp Fig 3).

### Aging biomarker genes selected by TimeFlies are differentially expressed in Drosophila Alzheimer’s models

To further assess the biological relevance of our model-selected genes, we sought to determine whether they are implicated in age-associated neurological disease. The Alzheimer’s Disease Fly Cell Atlas (ADFCA) was recently released [48], which includes whole-organism single-cell transcriptomics in Alzheimer’s Disease fly models. One model expresses the amyloid-beta 42 peptide (A®42) while the other expresses the wild-type human Tau (hTau) protein in the neurons of the brain [48]. We determined which genes were differentially expressed between AD genotypes (at the final stage of progression) and age-matched controls. We performed this analysis in a sex-specific manner for CNS neurons and sensory neurons, as they are the most abundant cell types in the ADFCA [48]. After obtaining the sets of differentially expressed genes (DEGs) for each sex and cell type, we compared them with our 50 most influential TimeFlies genes (determined by SHAP) for the corresponding sex/cell type. Many of the TimeFlies aging biomarker genes were either upregulated or downregulated in late-stage AD models (Fig. 4).

**Figure 4.**
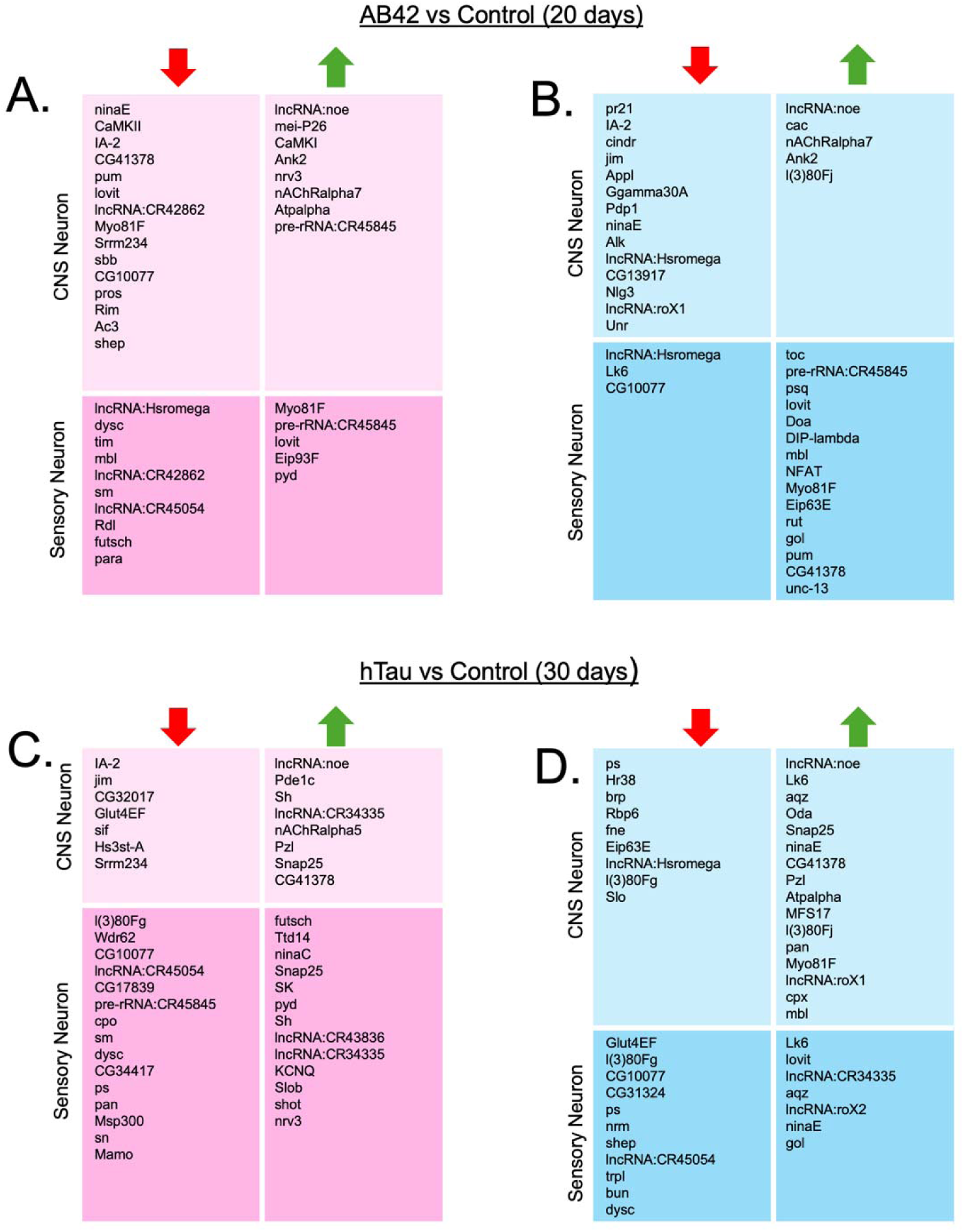
Genes that are both predictive aging genes from TimeFlies and differentially expressed in **A)** Aβ42 females, **B)** Aβ42 males, **C)** hTau females, **D)** hTau males. Differential expression analyses and comparison of gene sets were done in a sex and cell-type-specific manner, i.e., DEGs from male sensory neurons were compared with the predictive genes from the male sensory neuron TimeFlies clock. Cell types included in the analysis are CNS and sensory neurons. Left columns labeled with a downward-pointing red arrow contain genes downregulated in AD, while right columns labeled with an upward-pointing green arrow contain genes upregulated in AD.

Notably, the previously mentioned lncRNAs that were the most predictive of aging all appeared among the DEGs. *noe* is upregulated in both AD genotypes and in both sexes in both CNS and sensory neurons (Fig. 4A-D). *hsr*_ω_ is downregulated in A®42 female sensory neurons and hTau male CNS neurons, and both CNS and sensory neurons in A®42 males (Figure 4A-D). *roX1* is upregulated in A®42 male CNS neurons and hTau male CNS neurons, while *roX2* is upregulated in hTau male sensory neurons (Fig. 4B, D). *CR34335* is upregulated in hTau female sensory and CNS neurons, but only in sensory neurons in hTau males (Figure 4C, D). This analysis further solidifies the importance of these lncRNA genes to sex-specific cellular aging processes.

### Adult neuron depletion of CLAMP, an activator of the roX lncRNAs in males and a repressor in females, leads to a decline in lifespan in both males and females

The *roX* RNAs were the most significant hits from our TimeFlies aging clock. CLAMP is the primary regulator of the expression of the *roX* RNAs because it binds directly to the *roX* loci and tightly regulates their expression. CLAMP is required for regulating the levels of *roX* lncRNAs in both sexes because it increases levels of *roX* RNAs in males and represses *roX* RNAs in females [49, 50]. CLAMP has a stronger effect on *roX* RNA transcription in males versus females and males die earlier in development in the absence of CLAMP than females, but it is required for the viability of both sexes [50] (Fig. 5Ai). Therefore, activation of the *roX* in males and repression in females are both essential for normal development. However, nothing was known about how dysregulation of the *roX* RNAs by depleting CLAMP alters lifespan.

**Figure 5.**
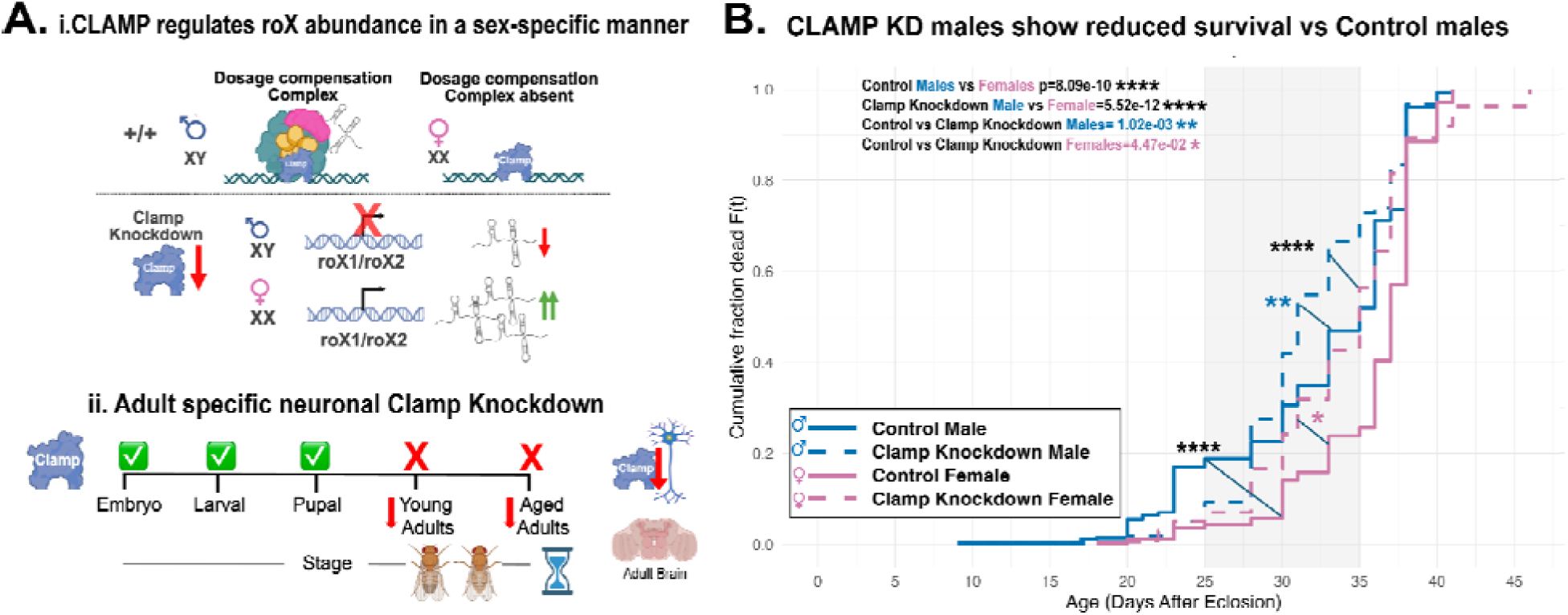
*Adult-specific neuronal CLAMP knockdown reduces male lifespan more significantly than female lifespan.* (**A**) Schematic of the experimental design showing pan-neuronal knockdown of CLAMP in adult flies. CLAMP normally binds GA-rich MSL recognition elements near *roX1* and *roX2* loci to recruit the MSL complex and mediate X-chromosome dosage compensation in males. In females, CLAMP maintains the repression of the roX RNAs. Therefore, CLAMP is required to regulate *roX* RNAs in both males and females. (**B**) Lifespan analysis revealed a significant reduction in male survival following CLAMP knockdown (Control vs *clampRNAi* females, *n* = 277 Clamp KD males, 326 Clamp KD females, 340 control males, 272 control females), while females were also affected (4.47e-02) but not as significantly as males (p=1.02e-03). Pan-neuronal CLAMP knockdown shortened lifespan in both sexes, with males more affected than females, suggesting that a precise balance of roX levels is essential for a normal lifespan just as it is during development.

We demonstrate that CLAMP depletion in adult neurons significantly shortens male lifespan (p=1.02e-03), while females were also affected, although not as significantly (4.47e-02) (Fig. 5A,–B). Therefore, our data suggests that highly regulated expression of the *roX* RNAs is also required for a normal lifespan in addition to normal development. Together, these results suggest that CLAMP-dependent transcriptional balance contributes to male-biased aging trajectories in the fly brain, and that perturbing neuronal dosage compensation accelerates age-associated decline through mechanisms overlapping with neurodegenerative pathways.

## Discussion

We have developed an aging clock based on explainable deep neural networks that classifies *Drosophila melanogaster* age at single-cell resolution with high accuracy. We have used the Aging Fly Cell Atlas [19], a diverse atlas of gene expression dynamics in the fly head at four time points across the lifespan. Our clock, TimeFlies, requires no feature engineering prior to training, unlike its predecessors. Importantly, it generalizes to all cell types despite the significant sparsity and high variability of single-cell RNA-seq data. While our study focuses on *Drosophila melanogaster* head tissue, this framework may be used for any single-cell RNA-seq aging atlas dataset, regardless of tissue or species, upon retraining with the dataset of interest.

Following the training and testing of our clock, we performed a feature explanation using Shapley values to discover potential transcriptomic signatures of aging. We found that long noncoding RNAs were enriched in TimeFlies feature explanations, consistent with previous studies that suggest an important role for lncRNAs in aging [25, 26, 27]. lncRNA:*roX1* and lncRNA:*roX2*, noncoding RNAs on the X chromosome involved in the process of dosage compensation, were the top features across cell types. An additional top feature wa lncRNA:Hsromega (*hsr*_ω_), which is a developmentally essential noncoding RNA that organizes omega speckles in the nucleus. Two other top features were lncRNA:*noe* and lncRNA:*CR343355*, which have unknown functions and very limited associated literature. Explanation analysis of our model indicates that lncRNA-mediated gene regulation events may be significant to brain aging processes, calling for further investigation. These results, coupled with prior research which highlights aging-associated lncRNAs across diverse species from mammals to nematodes [26], make the case for inclusion of noncoding genes in transcriptomic aging clocks for all species.

Further feature explainability analysis revealed noteworthy differences in female-specific and male-specific clocks, implying that aging is highly sex-specific, even at single-cell resolution. The female clock was unable to generalize to male test data, and the male clock was unable to generalize to the female test data. We trained and tested sex-specific clocks for each cell type of interest; model explanations showed significant sex differences in clock genes in all tested cell types. CNS Neurons are the cell type with the largest difference in clock genes between males and females (Figure 3C). Upon performing GSEA, we discovered that the male CNS neuron clock genes were enriched for splicing-related pathways, which was not observed in the female CNS neuron clock. These results further call for the exploration of sex-specific splicing and its effects in *Drosophila* aging. Overall, our analysis highlights the need to develop aging clocks in a sex-specific manner to better inform our understanding of aging and intervention/rejuvenation research efforts.

It should be noted that, while SHAP scores are a mathematically robust framework for computing the influence of features on model predictions, a high SHAP value only indicates predictive importance within the given model. Features with high average SHAP value magnitudes may be interpreted as potential candidate longevity genes in this context. However, we cannot determine biological causality from SHAP values alone. Thus, we set out to test the findings of TimeFlies with *in vivo* biological validation.

The identification of the *roX* noncoding RNAs as top clock genes is especially interesting due to a similar finding in the mouse brain. snRNA-seq of the aging female mouse hypothalamus revealed age-differential expression of *Xist*, the mouse analog of *roX* genes. Furthermore, corresponding clocks based on that dataset identified *Xist* expression as a predictive factor in neuronal aging [32]. Given these results and our objective to test the biological relevance of the TimeFlies learned features, we sought to assess the role of dosage compensation in fruit fly aging. We found that adult-specific neuronal knockdown of the dosage compensation factor CLAMP, the primary regulator of the *roX* RNAs, specifically shortened male lifespan, indicating that disruption of CLAMP-dependent regulation of *roX* RNAs and the MSL complex accelerates male-biased aging in Drosophila.

These findings link predictions from single-cell transcriptomic data with organismal phenotypes, revealing that dosage compensation machinery may contribute to the sex-specific trajectories of *Drosophila* aging. The previous finding of *Xist* as a predictor of neuronal aging in mice [32], coupled with our results, implies that dosage compensation may have a role in aging across diverse species with sexually dimorphic lifespans. Our results emphasize the necessity to include sex- and cell-type specificity in aging clocks and demonstrate how explainable deep learning frameworks like TimeFlies can identify functionally relevant molecular regulators. Future work will dissect how lncRNAs such as *roX1* and *roX2* regulate chromatin organization and neuronal longevity, advancing our understanding of how dosage compensation and RNA-based regulation shape the aging brain.

## Methods

### Dataset

The Aging Fly Cell Atlas (AFCA) [19] is a publicly available dataset documenting the single-cell transcriptomic profiles of fruit flies at ages 5, 30, 50, and 70 days. It includes both fly head samples and body samples. Here, we focus on the head data, with the objective of better understanding the aging fruit fly brain. The fly head dataset contains 289981 cells across 16 broad cell types and 15992 genes. The AFCA includes a near-equal distribution of the male and female samples, unlike many other published aging atlases. This allows for the investigation of sex-specific aging dynamics. The authors of the AFCA have published their own aging clocks on the dataset. However, these clocks are trained on very specific cellular subtypes and do not generalize to the whole dataset, nor do they specifically address sex differences. The authors also performed feature interpretation on their clocks, but the analysis of these features was limited to ribosomal protein-coding genes, with very limited discussion of other relevant genes [19]. This leaves the door open for a new clock that generalizes across all cell types for the AFCA dataset and a comprehensive analysis of sex-differential transcriptomic patterns in aging, which are especially prevalent in the brain.

To ensure that our model was learning genuine biological signals rather than batch effects, we generated a batch-corrected dataset using scVI [51] (Supplementary Figure 4).

The input data is initially a sparse matrix, meaning a matrix format that does not explicitly store zero-valued data to conserve space and memory. Specifically, it is in a coordinate format (COO), consisting of the coordinates of the non-zero values. Prior to providing this data as input to TimeFlies, we converted this matrix to a dense matrix using the numpy [52] and scipy [53] libraries in Python.

### Model development and interpretation

To choose the framework for the TimeFlies aging clock, we tested several types of machine learning and deep learning models, namely, ElasticNet logistic regression classifier and RandomForest, both of which were implemented with the scikit-learn library [54], XGBoost, which was implemented with the xgboost library [55], and a simple multilayer perceptron (MLP) neural network, which was implemented with Tensorflow [56], and a convolutional neural network, which was implemented in Tensorflow as well [56]. The 1D CNN outperformed every model (Fig 6A). CNNs have been used for genomic applications to predict regulatory activity from sequential genomic data like DNA sequences, etc. [57–60], but to our knowledge, they have not been used for transcriptomic aging clocks. We selected a CNN-based model for TimeFlies architecture due to its comparatively high performance and efficiency (Fig. 6A).

**Figure 6.**
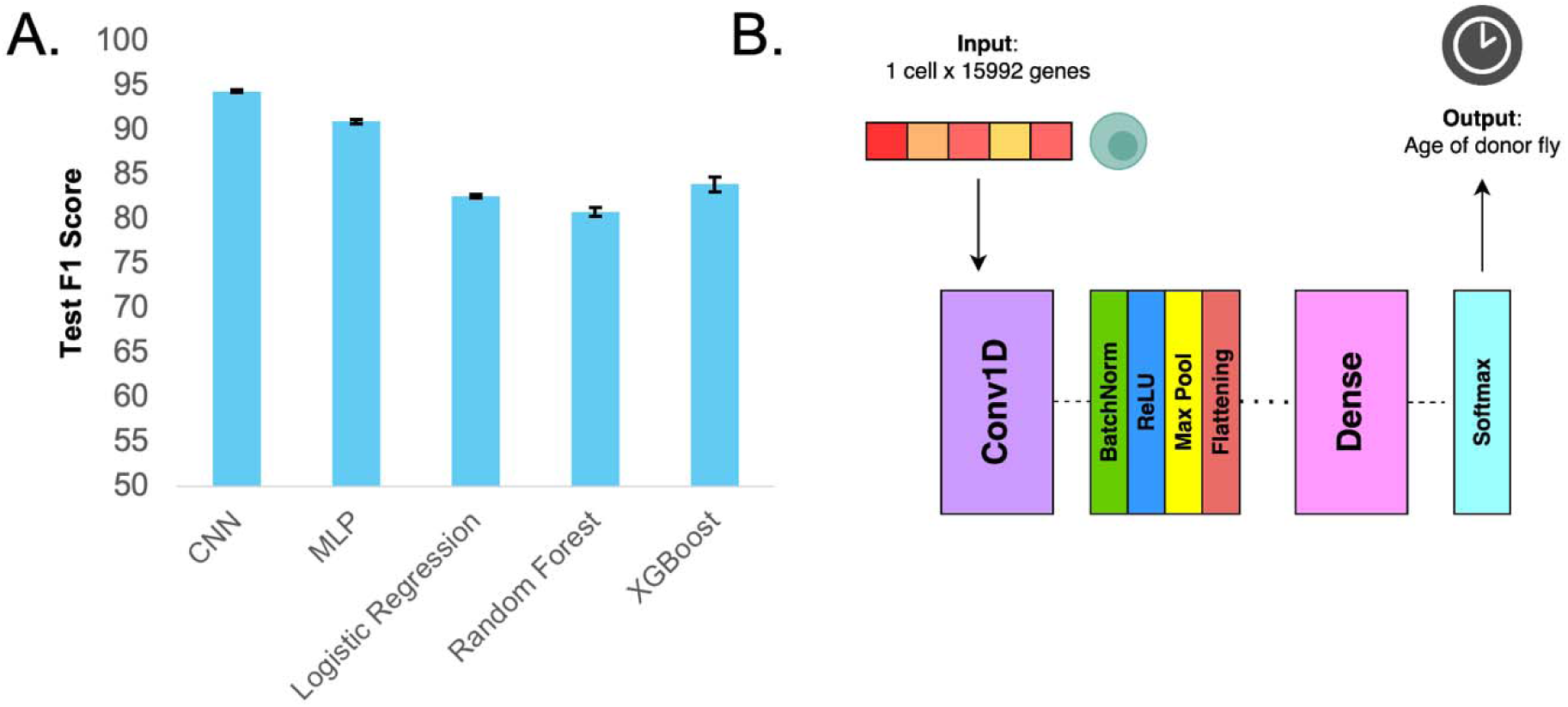
**A)** Comparison of performance on held-out test dataset across five different model types. Performance was tested across five random seeds, with error bars indicating standard deviation. **B)** Detailed model architecture of TimeFlies framework. TimeFlies consists of a 1D convolution layer and a dense layer (separated by batch normalization, nonlinear activation, and max pooling, and a flattening operation). Softmax activation is applied to the output, which is the age of the donor fly.

To optimize the architecture of the convolutional neural network, we tested three different combinations of convolutional and fully connected layers, the metrics of which are detailed in Supp. Tables 1-5. The TimeFlies aging clock (detailed in Fig. 6B) utilizes a convolutional neural network (CNN) consisting of 1D convolution blocks, pooling/flattening operations, and a dense layer.

Moreover, the features—genes—in the AFCA dataset were originally organized solely in alphabetical order without obvious spatial significance. However, shuffling the gene order with several different seeds has no significant impact on the TimeFlies performance (Supplementary Table 6) or model interpretation. Hence, we felt comfortable proceeding with the 1D CNN framework for the final architecture of TimeFlies.

TimeFlies is implemented in Python with the Tensorflow library [56]. We do not perform any feature selection of genes and input the transcriptome-wide gene expression profile. An input sample is a vector of gene expression for a single cell. All the samples are split into training, validation, and test sets of 80%, 10%, and 10%, respectively. Due to the high dimensionality of the dataset, GPU acceleration was used to speed up training time for TimeFlies and its benchmark models. Feature explanation analysis of TimeFlies was performed by obtaining Shapley values from GradientExplainer [61] and observing the features ranked highest. GradientExplainer is a method of approximating Shapley scores designed specifically for deep neural networks; it works by calculating the model’s output gradients with respect to each feature along a path from a baseline input to the actual input sample. The resulting Shapley scores are an average of these gradients over multiple baseline-to-actual input paths [61]. Higher relative Shapley value magnitudes indicate that the gene is more influential in driving model predictions. Gene set enrichment analysis was performed in R with g:Profiler [62].

### Comparison with Alzheimer’s DEGs

The Alzheimer’s Disease Fly Cell Atlas (ADFCA) [48] is another publicly available dataset released by the same lab as the AFCA. It documents the single-cell transcriptomic profiles in two fly models of Alzheimer’s Disease (AD) along with age-matched controls. The fly head subset contains 360036 samples and 16219 genes. To perform the differential gene expression analysis, we used the Scanpy Python library [63] with the nonparametric Wilcoxon rank sum test. We performed this analysis for CNS neurons and sensory neurons in a sex-specific manner.

### Lifespan assay for Clamp Knockdown in adult neurons

We used the conditional temperature-sensitive *tub-GAL80ts* system to drive adult-specific neuronal knockdown of CLAMP in a uniform *w^1118^*genetic background to control for X-chromosome differences. Males carrying *w^1118^; UAS-clamp RNAi* (BL: 57163) were crossed with virgin females carrying *w^1118^; tub-GAL80ts; nSyb-GAL4*, which restricts expression to neurons and enables temporal control of induction. Crosses were maintained at 19°C and flipped every alternate day. Upon eclosion (day 0), F1 progeny (*w^1118^; tub-GAL80ts; nSyb-GAL4 > w^1118^; UAS-clamp RNAi; +/+*) were collected and allowed to mate for two days. Groups of 15 males or 15 females were then transferred to fresh food vials. Experimental flies were shifted to 29°C from day 2 post-eclosion to inactivate GAL80ts and induce *UAS-clamp RNAi* expression, while *w^1118^* controls (*w^1118^; tub-GAL80ts; nSyb-GAL4 > w^1118^; +/+*) were treated identically. Maintaining flies at 19°C during development ensured suppression of GAL4 activity, thereby eliminating developmental effects of CLAMP or dosage compensation complex (DCC) perturbation. The efficiency of the RNAi line that targets *clamp* has been validated [64]. The functionality of the GAL80ts system was validated using a UAS-GFP reporter, which confirmed the absence of GFP expression at 19°C and robust GFP induction at 29°C, verifying temperature-dependent control of GAL4 activity. Flies were transferred to fresh food every two days, and deaths were recorded until all flies had died. Lifespan assays were performed in three independent biological replicates per group, and survival data were analyzed using Kaplan–Meier survival [65] and cumulative fraction dead curves with log-rank tests in R.

## Data Availability

The Aging Fly Cell Atlas is accessible at https://hongjielilab.shinyapps.io/AFCA/. The Alzheimer’s Disease Fly Cell Atlas is accessible at https://hongjielilab.org/adfca/. Both atlases are downloadable in h5ad file format.

## Code Availability

All code is available on GitHub at https://github.com/rsinghlab/TimeFlies.

## References

1. López-Otín C, Blasco MA, Partridge L, Serrano M, Kroemer G. Hallmarks of aging: An expanding universe. Cell. 2023;186(2):243–278. doi:10.1016/j.cell.2022.11.001

2. Hannum G, Guinney J, Zhao L, et al. Genome-wide Methylation Profiles Reveal Quantitative Views of Human Aging Rates. Molecular cell. 2012;49(2):359. doi:10.1016/j.molcel.2012.10.016

3. Horvath S. DNA methylation age of human tissues and cell types. Genome Biology. 2013;14(10):3156. doi:10.1186/gb-2013-14-10-r115

4. Levine ME, Lu AT, Quach A, et al. An epigenetic biomarker of aging for lifespan and healthspan. Aging (Albany NY). 2018;10(4):573–591. doi:10.18632/aging.101414

5. Lu AT, Quach A, Wilson JG, et al. DNA methylation GrimAge strongly predicts lifespan and healthspan. Aging (Albany NY). 2019;11(2):303–327. doi:10.18632/aging.101684

6. Lu AT, Fei Z, Haghani A, et al. Universal DNA methylation age across mammalian tissues. Nat Aging. 2023;3(9):1144–1166. doi:10.1038/s43587-023-00462-6

7. Vanyushin BF, Nemirovsky LE, Klimenko VV, Vasiliev VK, Belozersky AN. The 5-Methylcytosine in DNA of Rats: Tissue and Age Specificity and the Changes Induced by Hydrocortisone and other Agents. Gerontologia. 2009;19(3):138–152. doi:10.1159/000211967

8. Wilson VL, Smith RA, Ma S, Cutler RG. Genomic 5-methyldeoxycytidine decreases with age. J Biol Chem. 1987;262(21):9948–9951.

9. Romanov GA, Vanyushin BF. Methylation of reiterated sequences in mammalian DNAs. Effects of the tissue type, age, malignancy and hormonal induction. Biochim Biophys Acta. 1981;653(2):204–218. doi:10.1016/0005-2787(81)90156-8

10. Christensen BC, Houseman EA, Marsit CJ, et al. Aging and Environmental Exposures Alter Tissue-Specific DNA Methylation Dependent upon CpG Island Context. PLOS Genetics. 2009;5(8):e1000602. doi:10.1371/journal.pgen.1000602

11. Rutledge J, Oh H, Wyss-Coray T. Measuring biological age using omics data. Nat Rev Genet. 2022;23(12):715–727. doi:10.1038/s41576-022-00511-7

12. Peters MJ, Joehanes R, Pilling LC, et al. The transcriptional landscape of age in human peripheral blood. Nat Commun. 2015;6(1):8570. doi:10.1038/ncomms9570

13. Fleischer JG, Schulte R, Tsai HH, et al. Predicting age from the transcriptome of human dermal fibroblasts. Genome Biology. 2018;19:221. doi:10.1186/s13059-018-1599-6

14. Meyer DH, Schumacher B. BiT age: A transcriptome based aging clock near the theoretical limit of accuracy. Aging Cell. 2021;20(3):e13320. doi:10.1111/acel.13320

15. de Lima Camillo LP, Lapierre LR, Singh R. A pan-tissue DNA-methylation epigenetic clock based on deep learning. npj Aging. 2022;8(1):1–15. doi:10.1038/s41514-022-00085-y

16. Holzscheck N, Falckenhayn C, Söhle J, et al. Modeling transcriptomic age using knowledge-primed artificial neural networks. npj Aging Mech Dis. 2021;7(1):1–13. doi:10.1038/s41514-021-00068-5

17. Almanzar N, Antony J, Baghel AS, et al. A single-cell transcriptomic atlas characterizes ageing tissues in the mouse. Nature. 2020;583(7817):590–595. doi:10.1038/s41586-020-2496-1

18. Gao SM, Qi Y, Zhang Q, et al. Aging atlas reveals cell-type-specific effects of pro-longevity strategies. Nat Aging. 2024;4(7):998–1013. doi:10.1038/s43587-024-00631-1

19. Lu TC, Brbić M, Park YJ, et al. Aging Fly Cell Atlas identifies exhaustive aging features at cellular resolution. Science. 2023;380(6650):eadg0934. doi:10.1126/science.adg0934

20. Yu D, Li M, Linghu G, et al. CellBiAge: Improved single-cell age classification using data binarization. Cell Reports. 2023;42(12):113500. doi:10.1016/j.celrep.2023.113500

21. Mao S, Su J, Wang L, Bo X, Li C, Chen H. A transcriptome-based single-cell biological age model and resource for tissue-specific aging measures. doi:10.1101/gr.277491.122

22. Bronikowski AM, Meisel RP, Biga PR, et al. Sex-specific aging in animals: Perspective and future directions. Aging Cell. 2022;21(2):e13542. doi:10.1111/acel.13542

23. Dorkenwald S, Matsliah A, Sterling AR, et al. Neuronal wiring diagram of an adult brain. Nature. 2024;634(8032):124–138. doi:10.1038/s41586-024-07558-y

24. Schlegel P, Yin Y, Bates AS, et al. Whole-brain annotation and multi-connectome cell typing of Drosophila. Nature. 2024;634(8032):139–152. doi:10.1038/s41586-024-07686-5

25. Grammatikakis I, Panda AC, Abdelmohsen K, Gorospe M. Long noncoding RNAs (lncRNAs) and the molecular hallmarks of aging. Aging (Albany NY). 2014;6(12):992. doi:10.18632/aging.100710

26. Cai D, Han JDJ. Aging-associated lncRNAs are evolutionarily conserved and participate in NFκB signaling. Nat Aging. 2021;1(5):438–453. doi:10.1038/s43587-021-00056-0

27. Yang D, Lian T, Tu J, et al. LncRNA mediated regulation of aging pathways in Drosophila melanogaster during dietary restriction. Aging (Albany NY). 2016;8(9):2182. doi:10.18632/aging.101062

28. Paro PDR, Grossniklaus PDU, Santoro DR, Wutz PDA. Dosage Compensation Systems. In: Introduction to Epigenetics [Internet]. Springer; 2021. doi:10.1007/978-3-030-68670-3_4

29. Lentini A, Cheng H, Noble JC, et al. Elastic dosage compensation by X-chromosome upregulation. Nat Commun. 2022;13(1):1854. doi:10.1038/s41467-022-29414-1

30. Franke A, Baker BS. The rox1 and rox2 RNAs Are Essential Components of the Compensasome, which Mediates Dosage Compensation in Drosophila. Molecular Cell. 1999;4(1):117–122. doi:10.1016/S1097-2765(00)80193-8

31. Lucchesi JC, Kuroda MI. Dosage Compensation in Drosophila. Cold Spring Harb Perspect Biol. 2015;7(5):a019398. doi:10.1101/cshperspect.a019398

32. Meller VH, Rattner BP. The roX genes encode redundant male-specific lethal transcripts required for targeting of the MSL complex. The EMBO Journal. 2002;21(5):1084. doi:10.1093/emboj/21.5.1084

33. Hajdarovic KH, Yu D, Hassell LA, et al. Single-cell analysis of the aging female mouse hypothalamus. Nat Aging. 2022;2(7):662–678. doi:10.1038/s43587-022-00246-4

34. Lakhotia, S. C. Forty years of the 93D puff of Drosophila melanogaster. J Biosci 36, 399–423 (2011).

35. Singh, A. K. & Lakhotia, S. C. Dynamics of hnRNPs and omega speckles in normal and heat shocked live cell nuclei of Drosophila melanogaster. Chromosoma 124, 367–383 (2015).

36. Mallik, M. & Lakhotia, S. C. Pleiotropic consequences of misexpression of the developmentally active and stress-inducible non-coding hsrω gene in Drosophila. J Biosci 36, 265–280 (2011).

37. Singh, A. K. Hsrω and Other lncRNAs in Neuronal Functions and Disorders in Drosophila. Life 13, 17 (2023).

38. Davie, K. et al. A Single-Cell Transcriptome Atlas of the Aging Drosophila Brain. Cell 174, 982–998.e20 (2018).

39. Kim B, Shortridge RD, Seong C, Oh Y, Baek K, Yoon J. Molecular Characterization of a Novel *Drosophila* Gene Which Is Expressed in the Central Nervous System. Molecules and Cells. 1998;8(6):750–757. doi:10.1016/S1016-8478(23)13493-5

40. Perez G, Barber GP, Benet-Pages A, et al. The UCSC Genome Browser database: 2025 update. Nucleic Acids Res. Published online October 26, 2024:gkae974. doi:10.1093/nar/gkae974

41. Johnson K, Knust E, Skaer H. *bloated tubules (blot)* Encodes a *Drosophila* Member of the Neurotransmitter Transporter Family Required for Organisation of the Apical Cytocortex. Developmental Biology. 1999;212(2):440–454. doi:10.1006/dbio.1999.9351

42. Brown JB, Boley N, Eisman R, et al. Diversity and dynamics of the Drosophila transcriptome. Nature. 2014;512(7515):393–399. doi:10.1038/nature12962

43. Carrillo, R. A. et al. Control of Synaptic Connectivity by a Network of Drosophila IgSF Cell Surface Proteins. Cell 163, 1770–1782 (2015).

44. Camilleri-Robles, C., et al. Long non-coding RNAs involved in Drosophila development and regeneration. NAR Genom Bioinform 6, lqae091 (2024).

45. Bhadra, M., Howell, P., Dutta, S., Heintz, C. & Mair, W. B. Alternative splicing in aging and longevity. Hum Genet 139, 357–369 (2020).

46. Deschênes, M. & Chabot, B. The emerging role of alternative splicing in senescence and aging. Aging Cell 16, 918–933 (2017).

47. Ray, M. et al. Sex-specific splicing occurs genome-wide during early Drosophila embryogenesis. eLife 12, e87865 (2023).

48. Park, Y.-J. et al. Distinct systemic impacts of Aβ42 and Tau revealed by whole-organism snRNA-seq. Neuron 113, 2065–2082.e8 (2025).

49. Soruco, M. M. L. et al. The CLAMP protein links the MSL complex to the X chromosome during Drosophila dosage compensation. Genes Dev. 27, 1551–1556 (2013).

50. Urban, J. et al. Enhanced chromatin accessibility of the dosage compensated Drosophila male X-chromosome requires the CLAMP zinc finger protein. PLoS One 12, e0186855 (2017).

51. Gayoso, A. et al. A Python library for probabilistic analysis of single-cell omics data. Nat Biotechnol 40, 163–166 (2022).

52. Harris CR, Millman KJ, van der Walt SJ, et al. Array programming with NumPy. Nature. 2020;585(7825):357–362. doi:10.1038/s41586-020-2649-2

53. Virtanen P, Gommers R, Oliphant TE, et al. SciPy 1.0: fundamental algorithms for scientific computing in Python. Nat Methods. 2020;17(3):261–272. doi:10.1038/s41592-019-0686-2

54. Buitinck L, Louppe G, Blondel M, et al. API design for machine learning software: experiences from the scikit-learn project. Published online September 1, 2013. doi:10.48550/arXiv.1309.0238

55. Chen T, Guestrin C. XGBoost: A Scalable Tree Boosting System. Published online June 10, 2016. doi:10.48550/arXiv.1603.02754

56. Abadi M, Agarwal A, Barham P, et al. TensorFlow: Large-Scale Machine Learning on Heterogeneous Distributed Systems. Published online March 16, 2016. doi:10.48550/arXiv.1603.04467

57. Kelley DR, Snoek J, Rinn JL. Basset: learning the regulatory code of the accessible genome with deep convolutional neural networks. Genome Res. 2016;26(7):990–999. doi:10.1101/gr.200535.115

58. Singh R, Lanchantin J, Robins G, Qi Y. DeepChrome: deep-learning for predicting gene expression from histone modifications. Bioinformatics. 2016;32(17):i639–i648. doi:10.1093/bioinformatics/btw427

59. Kelley DR, Reshef YA, Bileschi M, Belanger D, McLean CY, Snoek J. Sequential regulatory activity prediction across chromosomes with convolutional neural networks. Genome Research. 2018;28(5):739. doi:10.1101/gr.227819.117

60. Alipanahi B, Delong A, Weirauch MT, Frey BJ. Predicting the sequence specificities of DNA- and RNA-binding proteins by deep learning. Nat Biotechnol. 2015;33(8):831–838. doi:10.1038/nbt.3300

61. Lundberg S, Lee SI. A Unified Approach to Interpreting Model Predictions. Published online November 25, 2017. doi:10.48550/arXiv.1705.07874

62. Kolberg L, Raudvere U, Kuzmin I, Adler P, Vilo J, Peterson H. g:Profiler—interoperable web service for functional enrichment analysis and gene identifier mapping (2023 update). Nucleic Acids Research. 2023;51(W1):W207–W212. doi:10.1093/nar/gkad347

63. Wolf, F. A., Angerer, P. & Theis, F. J. SCANPY: large-scale single-cell gene expression data analysis. Genome Biology 19, 15 (2018).

64. Kentro, J. A., et al. Conserved transcription factors coordinate synaptic gene expression through repression. bioRxiv 2024.10.30.621128 (2025) doi:10.1101/2024.10.30.621128.

65. Kaplan EL, Meier P. Nonparametric Estimation from Incomplete Observations. Journal of the American Statistical Association. 1958;53(282):457–481. doi:10.1080/01621459.1958.10501452

