## Supplementary Figures for "An snRNA-seq aging clock for the fruit fly head sheds light on sex-biased aging"

**Accuracy**

| CNN Type | **Seed: 42** | **Seed: 11** | **Seed: 1** | **Seed: 100** | **Seed: 999** |
|---|---|---|---|---|---|
| 1conv1fc | 94.1539% | 94.5168% | 94.5097% | 94.5524% | 93.9937% |
| 2conv2fc | 94.8975% | 94.7374% | 94.8121% | 94.3104% | 95.0185% |
| 3conv3fc | 94.8762% | 94.6093% | 94.6698% | 94.3069% | 94.9509% |

**Precision**

| CNN Type | **Seed: 42** | **Seed: 11** | **Seed: 1** | **Seed: 100** | **Seed: 999** |
|---|---|---|---|---|---|
| 1conv1fc | 93.4734% | 93.9497% | 93.9331% | 94.1297% | 93.7041% |
| 2conv2fc | 94.4830% | 94.2952% | 94.2918% | 93.7175% | 94.5789% |
| 3conv3fc | 94.3660% | 94.2390% | 94.1532% | 93.7161% | 94.4924% |

**Recall**

| CNN Type | **Seed: 42** | **Seed: 11** | **Seed: 1** | **Seed: 100** | **Seed: 999** |
|---|---|---|---|---|---|
| 1conv1fc | 93.7586% | 94.0209% | 94.0798% | 93.9768% | 93.3299% |
| 2conv2fc | 94.3633% | 94.2077% | 94.3493% | 93.8613% | 94.5425% |
| 3conv3fc | 94.4301% | 93.9631% | 94.2518% | 93.8720% | 94.6727% |

**F1**

| CNN Type | **Seed: 42** | **Seed: 11** | **Seed: 1** | **Seed: 100** | **Seed: 999** |
|---|---|---|---|---|---|
| 1conv1fc | 93.5843% | 93.9825% | 93.9924% | 94.0449% | 93.4709% |
| 2conv2fc | 94.4133% | 94.2489% | 94.3132% | 93.7385% | 94.5597% |
| 3conv3fc | 94.3928% | 94.0958% | 94.1603% | 93.7492% | 94.4643% |

**AUC**

| CNN Type | **Seed: 42** | **Seed: 11** | **Seed: 1** | **Seed: 100** | **Seed: 999** |
|---|---|---|---|---|---|
| 1conv1fc | 99.4305% | 99.4409% | 99.4400% | 99.4597% | 99.4080% |
| 2conv2fc | 99.5386% | 99.4909% | 99.4939% | 99.4734% | 99.5219% |
| 3conv3fc | 99.5270% | 99.4881% | 99.5061% | 99.5123% | 99.5511% |

**Supplementary Tables 1-5.** Accuracy, Precision, Recall, F1 Score, and AUC metrics for 3 different tested CNN model architectures. 1conv1fc refers to one convolution layer followed by one fully-connected layer, 2conv2fc refers to 2 convolution layers followed by 2 fully-connected layers, and 3conv3fc refers to 3 convolution layers followed by 3 fully-connected layers.

We tested 3 different convolutional neural network (CNN) architectures (Supplementary Tables 1-5), varying the number of convolution layers and fully-connected layers. 1conv1fc refers to one convolution layer followed by one fully-connected layer, 2conv2fc refers to 2 convolution layers followed by 2 fully-connected layers, and 3conv3fc refers to 3 convolution layers followed by 3 fully-connected layers. We found that the 1conv1fc architecture slightly underperforms in comparison to 2conv2fc, but there was no significant difference between 2conv2fc and 3conv3fc. Thus, we selected the 2conv2fc architecture for TimeFlies.

| Shuffle Seed | Test Accuracy (%) | Test Precision (%) | Test Recall (%) | Test F1 (%) | Test AUC (%) |
|---|---|---|---|---|---|
| 42 | 94.35 | 93.62 | 94.09 | 93.80 | 99.47 |
| 11 | 94.52 | 94.17 | 93.93 | 94.01 | 99.51 |
| 1 | 94.59 | 93.96 | 94.22 | 94.06 | 99.49 |
| 100 | 94.83 | 94.33 | 94.38 | 94.35 | 99.54 |
| 999 | 94.57 | 94.12 | 94.01 | 94.06 | 99.47 |

**Supplementary Table 6.** Performance of the TimeFlies clock with five different randomly selected shuffle seeds, which shuffle gene order.

The TimeFlies clock was run five times with five randomly selected seeds for shuffling the order of the genes prior to training the model. Shuffling the order of the genes had no effect on model performance when measuring five metrics on the held-out test data.

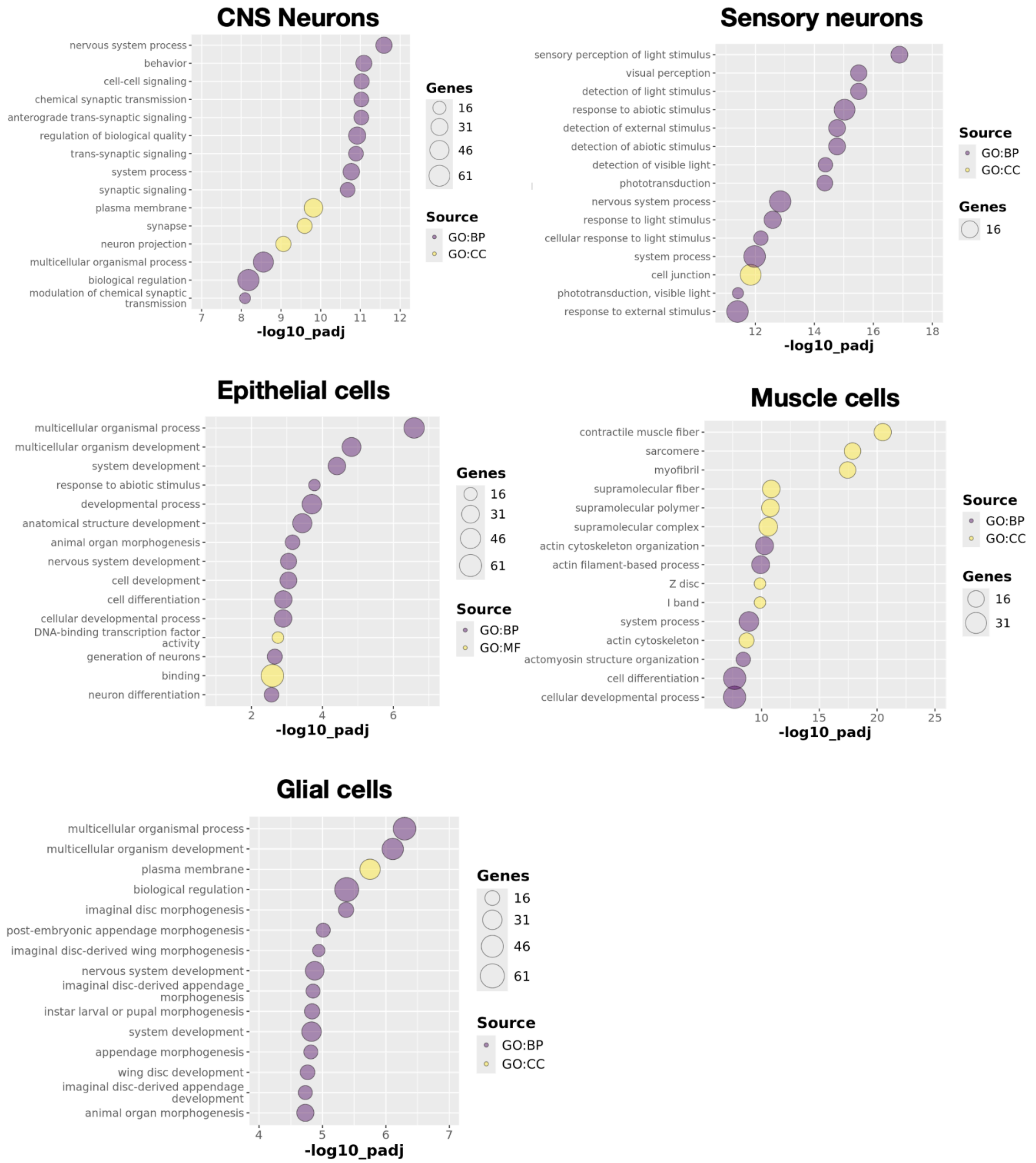


**Supplementary Figure 1.** Gene set enrichment analysis on the top 100 clock genes for each cell-type-specific clock. Purple bubbles indicate GO: biological process and yellow bubbles indicate GO: cellular component.

TimeFlies was trained on each broad cell type (CNS neurons, sensory neurons, epithelial cells, muscle cells, and glial cells) independently. The top fifty features for each clock–meaning, the genes with the highest Shapley score magnitudes–were evaluated as gene sets using g:Profiler in R. Gene ontology bubble plots were generated, with the more significantly enriched pathways/processes/cellular components at the top. The yellow bubbles correspond to the GO: cellular component category and the purple bubbles correspond to the GO: biological process category. In almost all cell types, there was a very clear association between the gene set enrichment analysis of the top clock genes and the known functions of the cell types; for glial cells, there is an enrichment of genes involved in disc/appendage/wing morphogenesis, but this is due to the pleiotropy of many of the glial clock genes. This validates that TimeFlies can learn cell-type-specific features while also having the ability to generalize across cell types in this dataset.

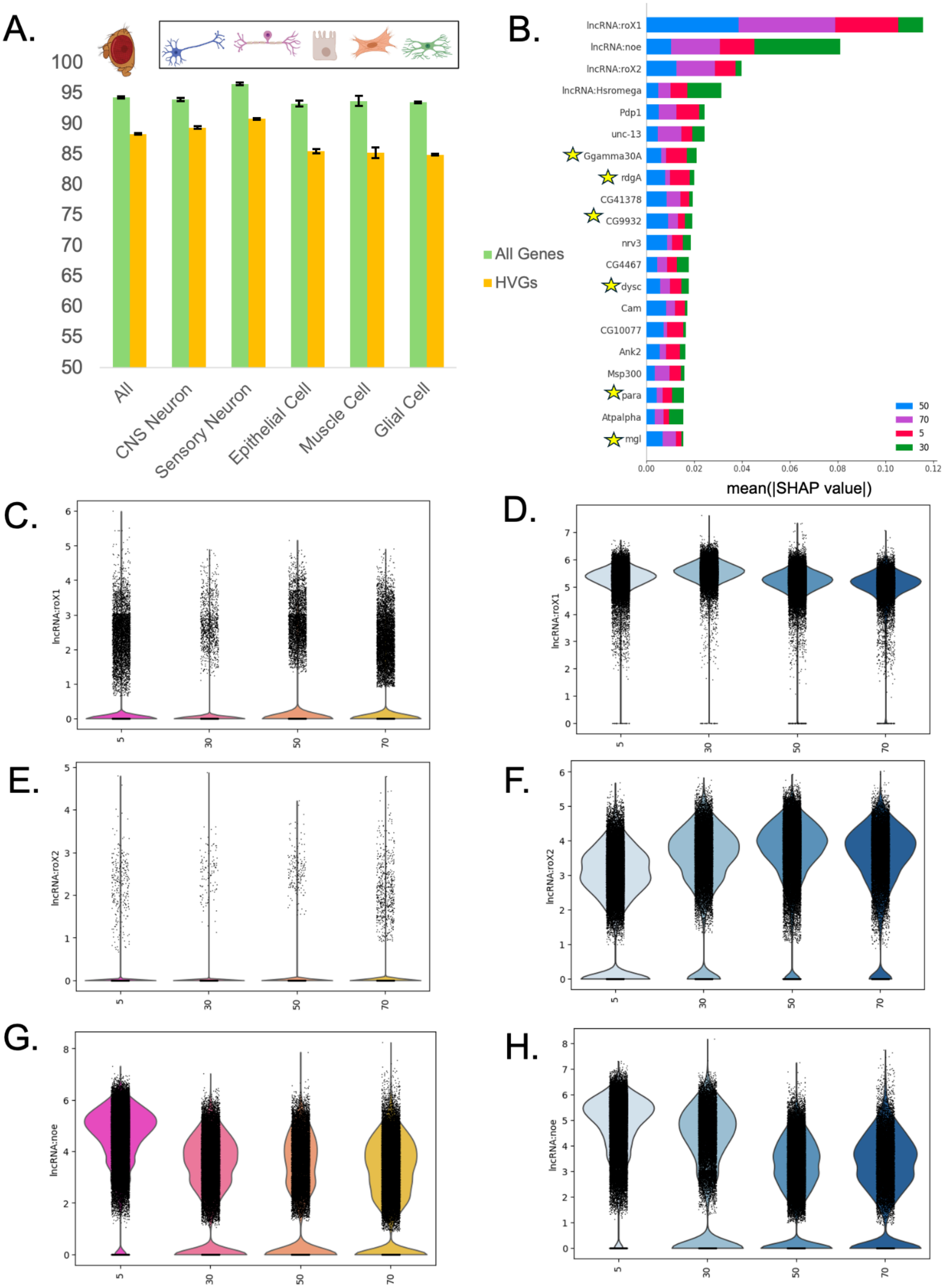


**Supplementary Figure 2.** A) Comparison of Test F1 Score with all genes as features (green) versus only the 5000 most highly variable genes as features (orange). B) In the top 20 features determined by Shapley analysis, those that are in the highly variable gene set are denoted with a star. C) Violin plot of lncRNA:*roX1* expression in females over time. D) Violin plot of lncRNA:*roX1* expression in males over time E) Violin plot of lncRNA:*roX2* expression in females over time. F) Violin plot of lncRNA:*roX2* expression in males over time. G) Violin plot of lncRNA:*noe* expression in females over time. H) Violin plot of lncRNA:*noe* expression in males over time.

In every cell type, using the whole set of features significantly outperformed HVG-based feature selection (Figure 2A). Moreover, when looking at the top 20 clock genes for the pan-cell-type model, only six appear in the set of 5000 most highly variable genes (Figure 2B). No significant linear expression pattern is observed with age in the top 3 pan-cell-type clock genes due to high variability of gene expression at each time point (Figure 2C-H).This shows that using the whole transcriptome rather than subsetting features boosts performance and that TimeFlies is able to capture complex mathematical patterns in the data that are often overlooked by more simple methods.

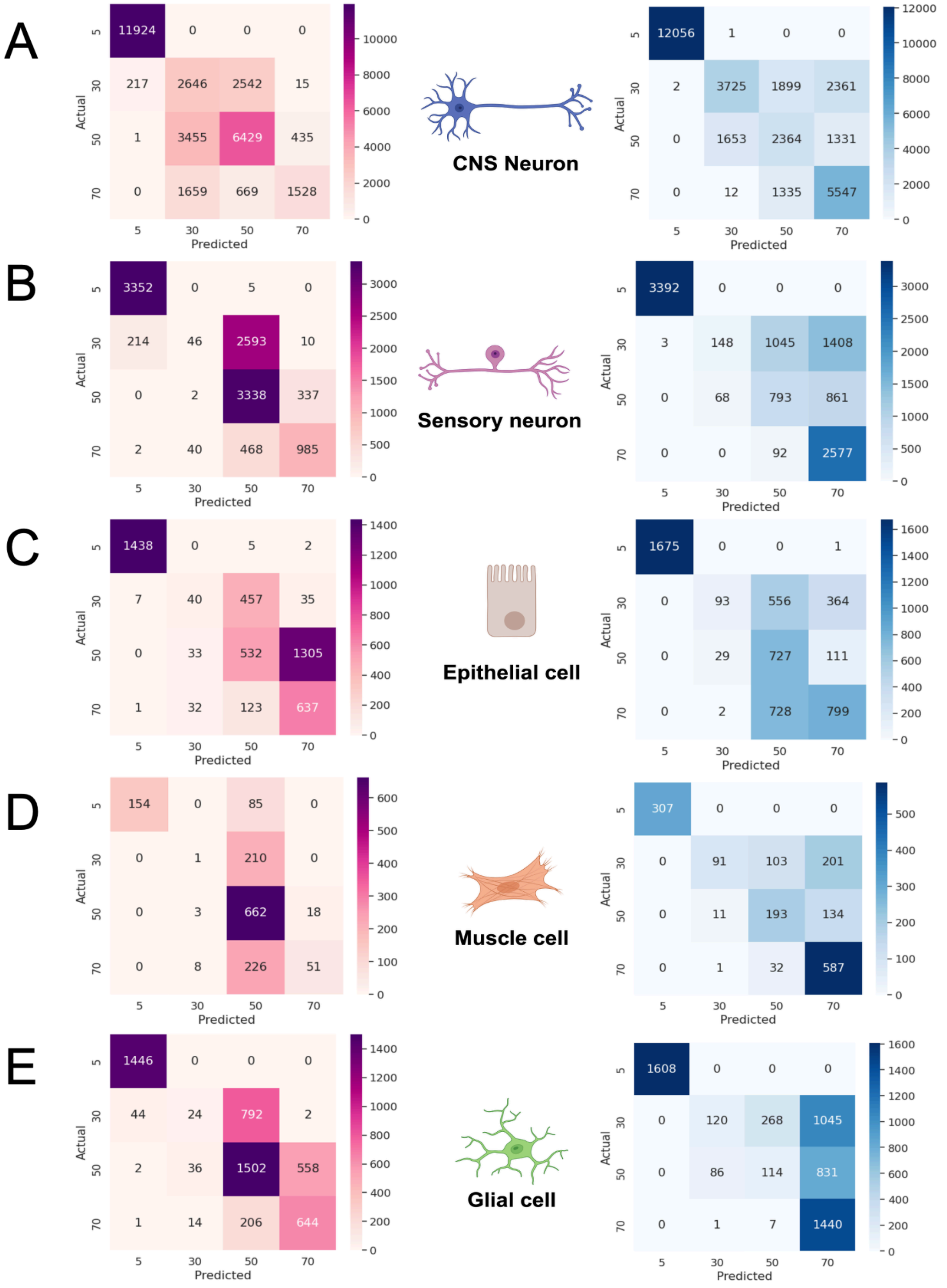


**Supplementary Figure 3. A)** Confusion matrix of female-trained CNS neuron clock tested on male data (left), confusion matrix of male-trained CNS neuron clock tested on female data (right). **B)** Confusion matrix of female-trained sensory neuron clock tested on male data (left), confusion matrix of male-trained sensory clock tested on female data (right). **C)** Confusion matrix of female-trained epithelial clock tested on male data (left), confusion matrix of male-trained epithelial clock tested on female data (right). **D)** Confusion matrix of female-trained muscle clock tested on male data (left), confusion matrix of male-trained muscle clock tested on female data (right). **E)** Confusion matrix of female-trained glial clock tested on male data (left), confusion matrix of male-trained glial clock tested on female data (right).

To determine the source of inaccuracy that results from training on one sex and testing on the other, we generated confusion matrices (Supplementary Fig 3). Notably, all cross-sex-tested clocks were able to distinguish 5-day-old cells with high accuracy. The male-trained clocks, apart from epithelial cell clocks, were also able to correctly classify 70-day-old female cells. The female-trained clocks were largely efficient in correctly classifying 50-day-old male cells, except for epithelial cells and CNS neurons.

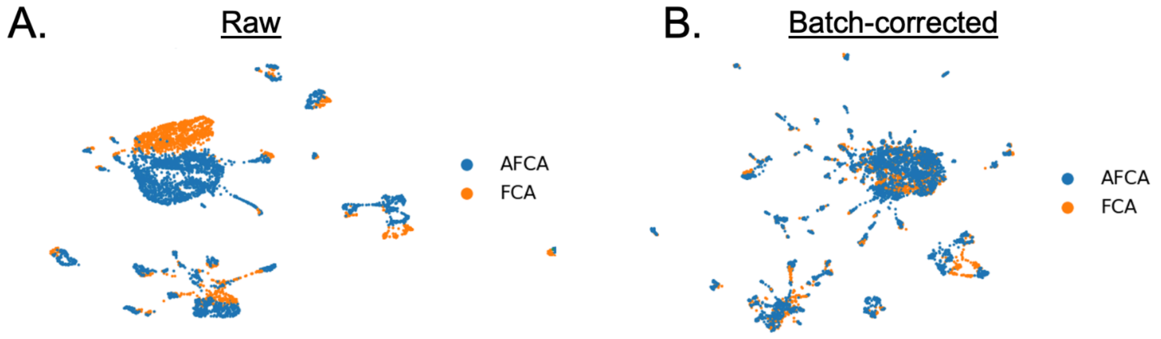


C.

| Data Type | Test Accuracy | Test F1 |
|---|---|---|
| Raw | 94.8762% | 94.3928% |
| Batch corrected | 94.67% | 94.11% |

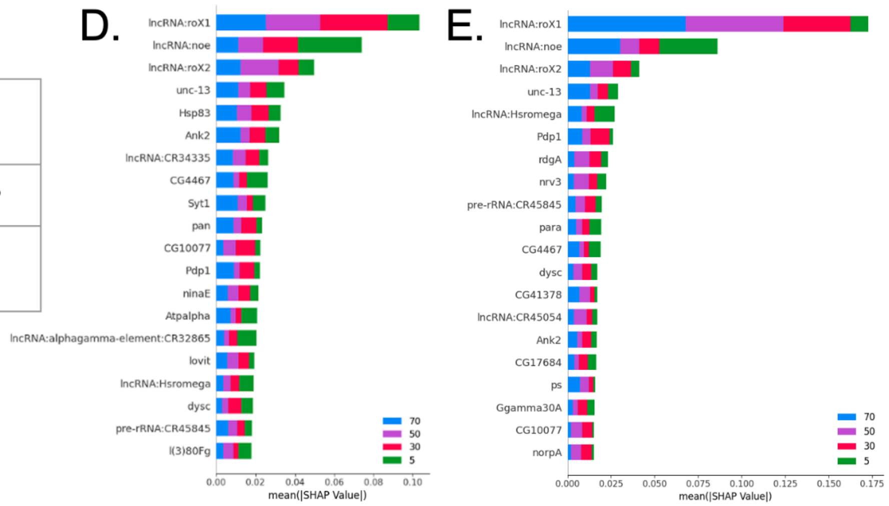


**Supplementary Figure 4.** A) UMAP of dataset before batch correction. B) UMAP of dataset after batch correction. C) Table of performance metrics of TimeFlies on raw and corrected datasets (both datasets use the whole transcriptome). D) SHAP summary plot of TimeFlies classification on raw dataset. E) SHAP summary plot of TimeFlies classification on the corrected dataset.

In our analysis, we addressed potential batch effects in the fly single-cell RNA-seq data using scVI for batch correction, aiming to mitigate any technical discrepancies that might obscure biological signals. We corrected for the “dataset” covariate, covariate which contained two categorical variables: AFCA and FCA, which are two separate sources of data that the authors integrated. After correction, we validated the integrity of the data by ensuring that the sample and gene orders remained consistent with the original dataset, confirming that no unintended alterations occurred during the batch correction process. To visually assess the impact of batch correction, we generated UMAP projections both before and after the correction. As illustrated in Supplementary Figure 4a-b, the UMAP plots reveal that the batch effect was eliminated.
